# Precision genetic cellular models identify therapies protective against endoplasmic reticulum stress

**DOI:** 10.1101/2020.06.03.132886

**Authors:** Irina V. Lebedeva, Michelle V. Wagner, Sunil Sahdeo, Yi-Fan Lu, Anuli Anyanwu-Ofili, Matthew B. Harms, Jehangir S. Wadia, Gunaretnam Rajagopal, Michael J. Boland, David B. Goldstein

## Abstract

Congenital disorders of glycosylation (CDG) and deglycosylation (CDDG) are a collection of rare pediatric disorders with symptoms that range from mild to life threatening. They typically affect multiple organ systems and usually present with neurological abnormalities including hypotonia, cognitive impairment, and intractable seizures. Several genes have been implicated in the thirty-six types of CDG, but currently *NGLY1* is the only known CDDG gene. A common biological mechanism among CDG types and in CDDG is endoplasmic reticulum (ER) stress. Here, we develop two isogenic human cellular models of CDG (*PMM2*, the most prevalent type of CDG, and *DPAGT1*) and of the only CDDG (*NGLY1*) in an effort to identify drugs that can alleviate ER stress. Systematic phenotyping identified elevated ER stress and autophagy levels among other cellular and morphological phenotypes in each of the cellular models. We screened a complex drug library for compounds able to correct aberrant morphological phenotypes in each of the models using an agnostic phenotypic cell painting assay based on >300 cellular features. The image-based screen identified multiple candidate compounds able to correct aberrant morphology, and we show a subset of these are able to correct cellular and molecular defects in each of the models. These results provide new directions for the treatment of rare diseases of glycosylation and deglycosylation and a framework for new drug screening paradigms for more common neurodegenerative diseases characterized by ER stress.

**Summary sentence:** Novel drug screening modality identifies compounds that correct aberrant molecular phenotypes in precision
cellular models of glycosylation defects.

## Introduction

Numerous diseases affecting both the central and peripheral nervous system involve elevated endoplasmic reticulum (ER) stress (*1, 2*). In particular, ER stress has been implicated in diseases including Parkinson’s disease, Alzheimer’s disease, and Amyotrophic Lateral Sclerosis (ALS) (*3*). This suggests that therapeutic agents that ameliorate the effects of ER stress could have benefits across a broad range of disorders. While screens to identify such agents in the context of complex neurodegenerative diseases are challenging to implement, the etiology of a number of monogenic diseases is in large part attributed to ER stress including the congenital disorders of glycosylation (CDG) and deglycosylation (CDDG) (*4-6*). Of particular interest, mutations in *PMM2*, the gene that encodes the cytosolic enzyme phosphomannomutase 2, result in the most common CDG, PMM2-CDG (*7*). Studies suggest that in PMM2*-*CDG, cells with weaker ER stress responses are more vulnerable to damage than cells with stronger ER stress responses (*8*). Moreover, mutations in *DPAGT1*, which encodes the target of the well-known ER stress inducer tunicamycin (*9*) result in another CDG with multisystemic phenotypes (*10, 11*).

Here we utilize a morphological profiling and screening paradigm to identify agents that protect against the cellular stresses resulting from CDG and CDDG causal mutations. We focus specifically on mutations in *PMM2* and *DPAGT1*, and in *NGLY1*, which causes the only reported CDDG (*6*). We used genetic engineering to generate clinically relevant CDG and CDDG genotypes in a karyotypically normal human cell line (hTERT RPE-1) in order to create cellular models amenable to mutation-specific phenotype identification. The hTERT RPE-1 line was selected to allow for morphology-based high content imaging screens to identify phenotypes that are consequences of ER stress. These CDG and CDDG cell lines were used in high-content small molecule screens to identify compounds that revert the imaging phenotypes caused by these mutations. Specifically, we selected 1049 annotated compounds for screening representing a broad chemical space and multiple target classes. In order to validate the performance of the screen, we selected 16 compounds that were ranked amongst the best at phenotype reversion in the screen (protective compounds) and 10 compounds that did not affect aberrant phenotypes (non-active negative control compounds). We then evaluated these compounds in assays designed to test how well they revert mutational phenotypes in our three cellular models.

## Results

### Establishment of precise human cellular models of CDG and CDDG

Genome editing was used to generate hTERT RPE-1 cell lines that mimic genotypes associated with CDG and CDDG (Table 1). All known CDDG patients possess complete loss of function of NGLY1 (*6*). We designed gRNAs to generate the recurrent *NGLY1* R401X missense variant, but after repeated attempts were unable to obtain the homozygous R401X genotype. Therefore, we screened clones for knock-out of *NGLY1* resulting from biallelic indel formation (*NGLY1*^-/-^). PMM2-CDG often results from compound heterozygous mutations that reduce enzymatic activity (*4, 12, 13*). Compound heterozygous *PMM2* lines were generated by monoallelic knock-in of the second most recurrent and very severe mutation (F119L) (*7, 14*), and then screening for an indel on the second allele (*PMM2*^F119L/-^). We generated *DPAGT1*^+/-^ lines by monoallelic knockout via indel formation. All genotypes were confirmed by Sanger sequencing (Figure 1A, Supplementary Figure 1, Supplementary Table 1).

**Table 1.**
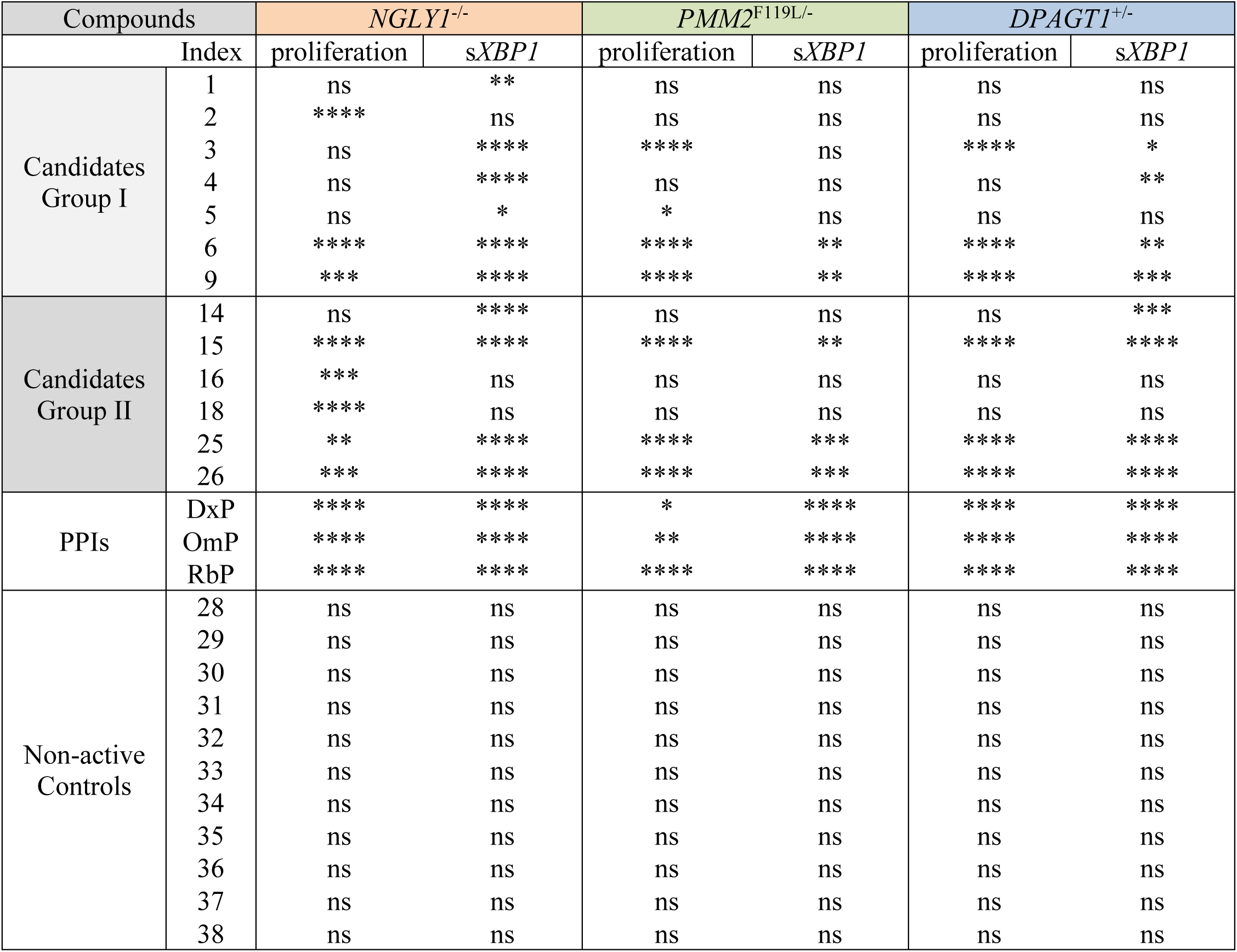
Correlation between the proliferation and s*XBP1* expression results in CDG and CDDG cell lines treated with candidate and non-active compounds, and PPIs. Statistical significance for s*XBP1* changes and proliferation rates changes. *, P<0.05, **, P<0.01, ***, P<0.001, ****, P<0.0001

| Compounds |  | *NGLY1*<sup>-/-</sup> |  | *PMM2*<sup>F119L/-</sup> |  | *DPAGT1*<sup>+/-</sup> |  |
| --- | --- | --- | --- | --- | --- | --- | --- |
| Index |  | proliferation | *sXBP1* | proliferation | *sXBP1* | proliferation | *sXBP1* |
| Candidates Group I | 1 | ns | ** | ns | ns | ns | ns |
|  | 2 | **** | ns | ns | ns | ns | ns |
|  | 3 | ns | **** | **** | ns | **** | * |
|  | 4 | ns | **** | ns | ns | ns | ** |
|  | 5 | ns | * | * | ns | ns | ns |
|  | 6 | **** | **** | **** | ** | **** | ** |
|  | 9 | *** | **** | **** | ** | **** | *** |
| Candidates Group II | 14 | ns | **** | ns | ns | ns | *** |
|  | 15 | **** | **** | **** | ** | **** | **** |
|  | 16 | *** | ns | ns | ns | ns | ns |
|  | 18 | **** | ns | ns | ns | ns | ns |
|  | 25 | ** | **** | **** | *** | **** | **** |
|  | 26 | *** | **** | **** | *** | **** | **** |
| PPIs | DxP | **** | **** | * | **** | **** | **** |
|  | OmP | **** | **** | ** | **** | **** | **** |
|  | RbP | **** | **** | **** | **** | **** | **** |
| Non-active Controls | 28 | ns | ns | ns | ns | ns | ns |
|  | 29 | ns | ns | ns | ns | ns | ns |
|  | 30 | ns | ns | ns | ns | ns | ns |
|  | 31 | ns | ns | ns | ns | ns | ns |
|  | 32 | ns | ns | ns | ns | ns | ns |
|  | 33 | ns | ns | ns | ns | ns | ns |
|  | 34 | ns | ns | ns | ns | ns | ns |
|  | 35 | ns | ns | ns | ns | ns | ns |
|  | 36 | ns | ns | ns | ns | ns | ns |
|  | 37 | ns | ns | ns | ns | ns | ns |
|  | 38 | ns | ns | ns | ns | ns | ns |

**Figure 1.**
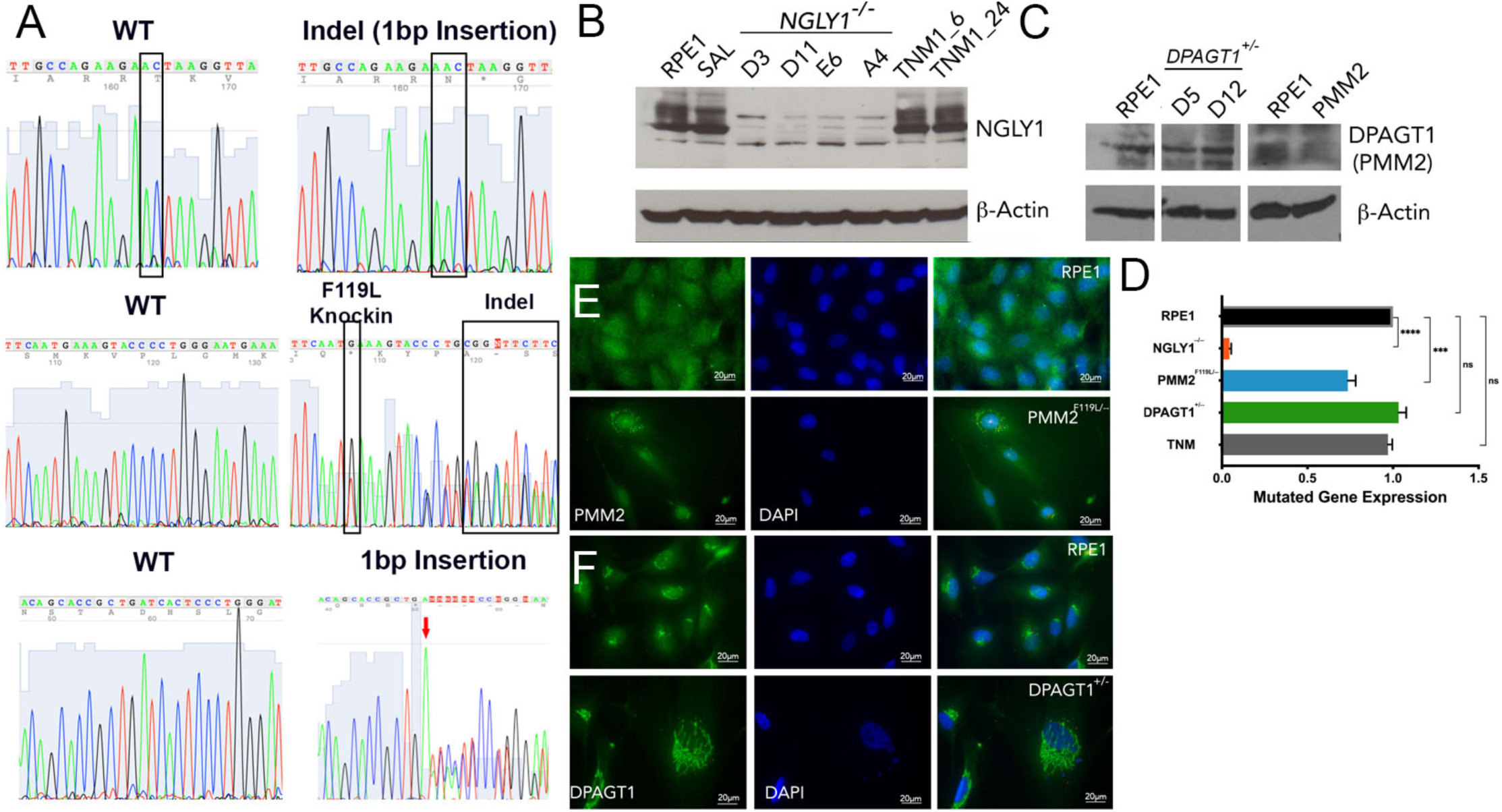
CDG and CDDG cellular models were validated by Sanger sequencing, immunoblot and immunofluorescence staining for target proteins. A, Electropherogram traces for parental RPE-1 cells, CDDG - *NGLY1*^-/-^ D11 (top), and CDG - *PMM2*^F119L/-^ A3 (middle) and *DPAGT1*^+/-^ D5 (bottom) lines. B and C, Immunoblot images for target proteins in RPE-1 and isogenic CDDG (B) and CDG (C) lines. RPE-1 cells treated with TNM (1µM for either 6 h or 24 h) were used as a positive control. D, Quantification of the target protein levels in the edited RPE-1 cells. Expression relative to levels in parental RPE-1 cells. ***, P<0.001, ****, P<0.0001. E and F, Representative immunofluorescence images for staining for PMM2 and DPAGT1 in parental RPE-1, CDG - *PMM2*^F119L/-^ A3, and CDG *DPAGT1*^+/-^ D5 lines

As expected, *NGLY1*^-/-^ lines do not express NGLY1 protein and the levels of PMM2 in *PMM2*^F119L/-^ were decreased by ∼50% (Figure 1B-C, Supplementary Figure 1). The expression level of DPAGT1 was different between the two *DPAGT1*^+/-^ clones analyzed despite confirmation of monoallelic disruption of *DPAGT1* (Figure 1C and Supplementary Figure 1). Clone DD5, however, consistently expressed ∼50% of DPAGT1 relative to parental cells. This clone was used in the high-content screens discussed below.

Consistent with published studies (*15, 16*), we found DPAGT1 localized to the perinuclear space, and PMM2 was diffuse throughout the cytosol and nucleus (Figure 1E-F, Supplementary Figure 1). Interestingly, PMM2^F119L^ was found in cytosolic puncta suggestive of protein aggregation (Figure 1E). We failed to detect NGLY1 by immunocytochemistry using multiple NGLY1 antibodies (not shown).

### CDG and CDDG lines exhibit elevated ER stress and autophagy responses

Although a common molecular feature of CDG is elevated levels of ER stress (*17*), systematic examination of ER stress in CDDG has not been performed. In order to establish the ER stress profiles of the cellular models, we first established a baseline in isogenic RPE-1 cells using a moderate concentration of the N-linked glycosylation inhibitor and ER stress inducer tunicamycin (*18, 19*) and the ER stress inhibitor salubrinal (*20*). We chose tunicamycin to activate the ER stress response because it is known to inhibit DPAGT1 (*9*). We examined ER stress using markers from each of the three recognized pathways of ER stress (Figure 2A): 1) detection of eIF2α Ser51 phosphorylation (peIF2α) and nuclear translocation of ATF4 (CHOP), 2) presence of spliced *XBP1* mRNA (s*XBP1*), and 3) cleavage of ATF6 (*19, 21*). As expected, tunicamycin treatment resulted in strong induction of peIF2α, elevated expression of s*XBP1*, and reduced levels of ATF6 (Figure 2C-E, TNM).

**Figure 2.**
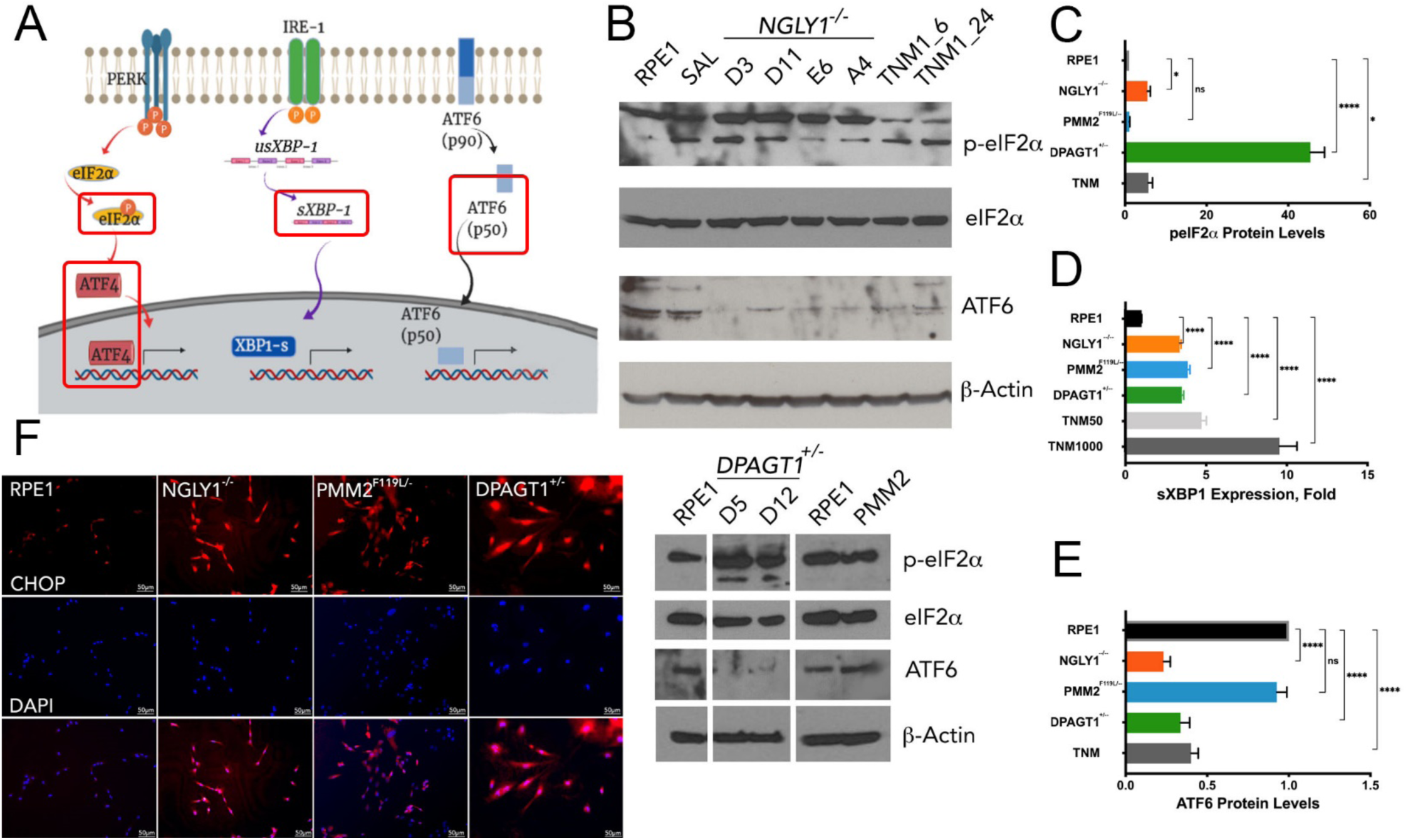
Cellular models of CDG and CDDG exhibit elevated ER stress responses. A, Schematic of the three major ER stress pathways (analyzed markers are framed). B, Immunoblot images for ER stress markers in RPE-1 and different clones of isogenic CDDG (top) and CDG (bottom) lines. RPE-1 cells treated with TNM (1µM for either 6 h or 24 h) and salubrinal (SAL, 50µM for 24 h) were used as negative and positive controls, respectively. C, Quantification of peIF2α levels relative to total eIF2α protein. D, Expression of spliced *XBP1* transcript. E, Quantification of ATF6 levels in parental and edited RPE-1 cells. Expression data in C-E are relative to levels in parental RPE-1 cells. *, P<0.05, **, P<0.01, ***, P<0.001, ****, P<0.0001. F, Nuclear localization of CHOP/ATF4 in parental and edited RPE-1 cells (scale bars are 50 µm).

CDG mutations are predicted to result in chronic ER stress (*17*). Indeed, all CGD and CDDG lines exhibited induction of increased ER stress relative to untreated RPE-1 (Figure 2B-E), but the distinct genotypes showed differential activation of the key ER stress response pathways. For example, *NGLY1*^-/-^ and *DPAGT1*^+/-^ lines exhibited activation of all three established ER stress pathways, whereas *PMM2*^F119L/-^ had only significant increases in *XBP1* splicing (Figure 2). In fact, the only pathway induced across all CDG and CDDG lines was splicing of *XBP1*. Low, but significant, levels of apoptosis were detected in *PMM2*^F119L/-^ and *NGLY1*^-/-^ but not in *DPAGT1*^+/-^ lines. Apoptosis levels in mutant cell lines were much lower as would expected from high ER stress (Supplementary Figure 2B-C) further suggesting that CDG and CDDG lines exhibit lower chronic ER stress responses.

Autophagy is known to play an important role in the response to ER stress and is it seen as a marker of chronic ER stress (*22*). Significant upregulation of autophagy was detected in all mutant cell lines using a cationic amphiphilic tracer dye that labels autophagic vacuoles (*23*) by both fluorescent microscopy (Figure 3A) and flow cytometry (Figure 3B). Autophagy induction was also seen with markers of early (p62/SQSTM1) and late (LAMP1) stages of autophagy followed by fluorescent microscopy (Figure 3C-E and Supplementary Figure 3).

**Figure 3.**
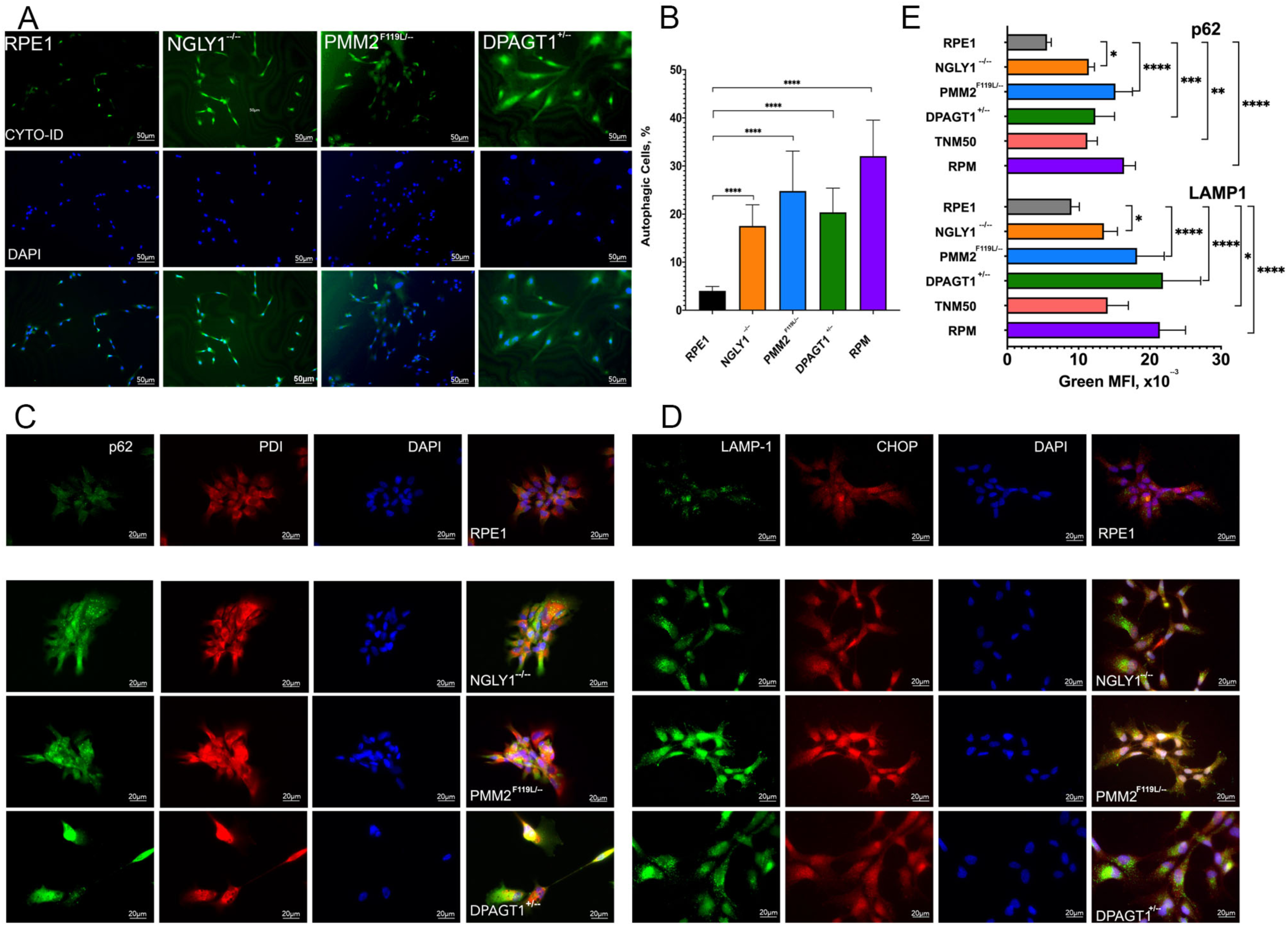
CDG and CDDG lines exhibit elevated autophagy levels. Cells were stained using CYTO-ID Autophagy Detection Kit and observed by (A) fluorescent microscopy; scale bar, 50 µm, and (B) analyzed by flow cytometry. C and D, Representative immunofluorescence images of parental RPE-1 cells and CDDG and CDG cell lines stained with antibodies against (C) p62/SQSTM1 or (D) LAMP1 in combination with anti-CHOP or anti-PDI antibodies, respectively. Scale bar, 20 µm. E, Quantification of p62/SQSTM1 and LAMP1 staining by flow cytometry. *, P<0.05, **, P<0.01, ***, P<0.001, ****, P<0.0001.

### CDG and CDDG lines exhibit distinctive morphological phenotypes and proliferation defects

CDG and CDDG lines were characterized for phenotypes useful for high-content imaging screens. All mutant cell lines exhibited a flat, extended morphology (Figure 4A) reminiscent of cellular senescence that was not seen in the isogenic parental line. Indeed, β-galactosidase staining confirmed various levels of senescence among the mutant cell lines (Figure 4B, Supplementary Figure 2A). All lines demonstrated slower proliferation compared to the isogenic RPE-1 line (Figure 4C). *DPAGT1*^+/-^ lines exhibited the slowest proliferation rates and were comparable to those observed in the parental line when subjected to chronic ER stress from low concentration tunicamycin exposure (Figure 4C).

**Figure 4.**
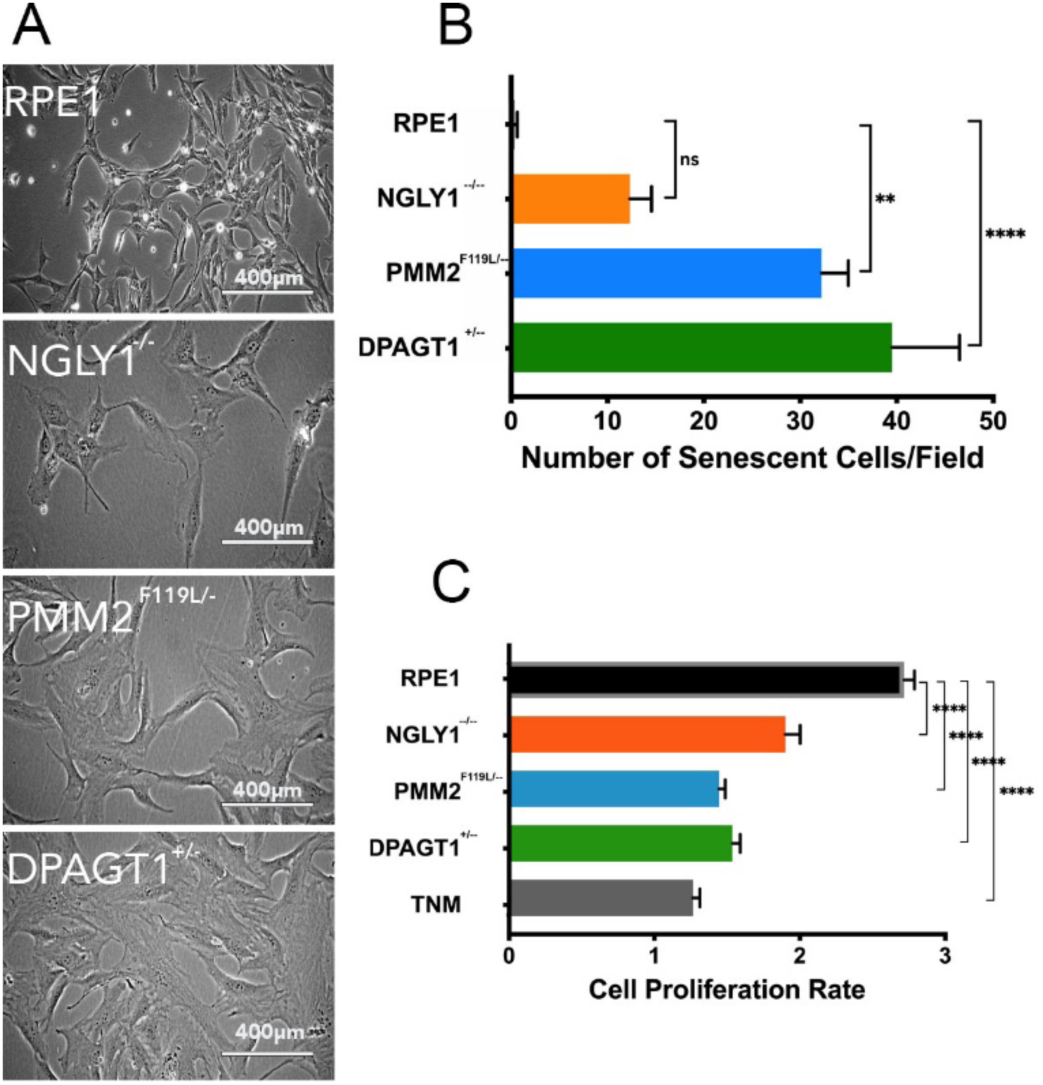
CDG and CDDG lines exhibit line-dependent levels of senescence and reduced proliferation. Isogenic RPE-1 and edited cell lines were seeded in tissue culture plates, cultured for several days, and phase contrast images were taken. Separate set of samples were fixed and stained for β-Galactosidase, an indicator of senescence. For proliferation assay, cells were seeded in triplicates in 96 well plates, stained every other day with MTT and OD_590_ was measured. A, Representative phase contrast images of cellular morphology. Scale bars, 400 µm. B, Quantification of senescence levels as indicated by β-galactosidase staining. C, Quantification of cellular proliferation rates for CDG and CDDG lines relative to parental RPE-1. To define cell proliferation rate, ratio of OD_590_ at 72 h to OD_590_ at 24 h post seeding was calculated. *, P<0.05, **, P<0,01, ****, P<0.0001

### Primary drug screen identifies compounds able to reverse CDG and CDDG cellular morphology phenotypes

Our drug screening platform takes advantage of the distinctive cellular phenotypes that result from the CDG and CDDG mutations. In order to identify compounds able to correct aberrant morphological phenotypes in the mutant lines, we utilized a “cell painting” phenotypic assay (*24, 25*) based on stains for mitochondria, the actin cytoskeleton, endoplasmic reticulum, and nuclei (Figure 5A). Machine learning algorithms were trained on acquired images of RPE-1 cells, and more than 300 cellular features such as fluorescence intensity, presence and numbers of puncta, texture, and cellular shape and geometry were extracted and analyzed. Functional testing and validation of the cell painting assay was performed on CDG and CDDG cell lines (Figure 5B). Importantly, hierarchical clustering and principal component analyses clearly distinguished mutant cells from each other and from parental RPE-1 cells (Figure 5C and D). This demonstrates that there are distinct phenotypic, morphological changes that occur as a consequence of the CDG or CDDG mutation in each of the clones.

**Figure 5.**
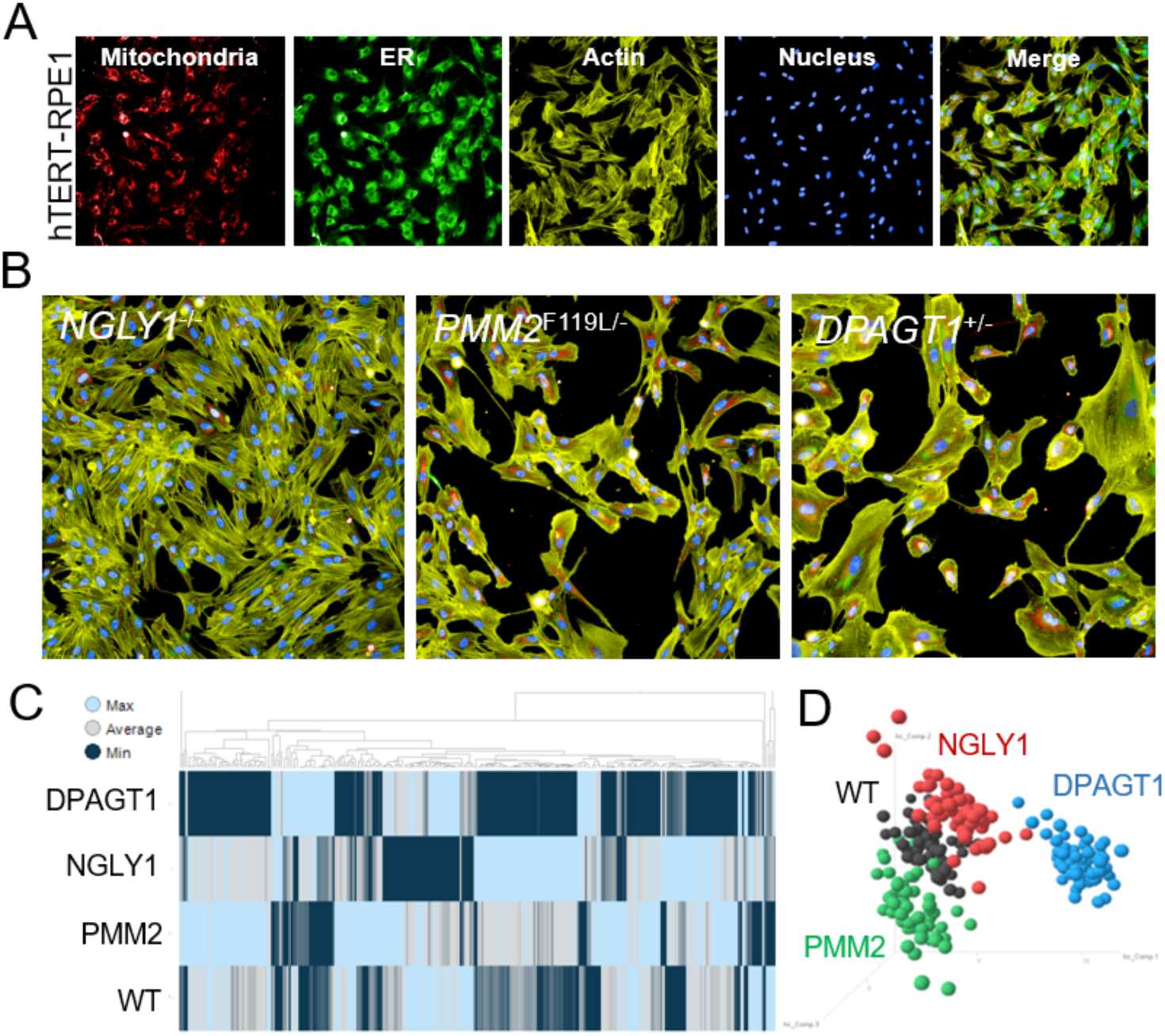
Cell painting assay developed for high-throughput cellular morphology screening identifies line-specific morphological characteristics. A, Cell painting images of parental RPE-1 cells. Cells were plated in 384-well tissue culture plate and stained with MitoTracker Red (mitochondrial stain), ConcanavalinA-488 (ER stain), phalloidin-547 (actin stain), and DAPI (nuclei) as described in the Methods section. B, Representative cell painting images of CDDG and CDG lines. C and D, Hierarchical clustering and principal component analyses of extracted morphological features distinguishes CDG and CDDG cell lines from parental RPE-1 cells

We screened 1,049 annotated compounds representing a broad chemical space and multiple target classes on CDG and CDDG cell lines (Figure 6A). The compound library was assembled with publicly available compounds that have known biological activities as well as Janssen proprietary compounds that have evidence of bioactivity compiled from multiple internal data sets. *NGLY1*^*-/-*^, *PMM2*^*F119L/-*^ *and DPAGT1*^*+/-*^ lines were treated for 24 hours with 10 µM of each compound. Parental RPE-1 cells treated with vehicle (DMSO) served as a positive control while vehicle-treated CDG and CDDG lines served as negative controls (Figure 6B). Post-treatment, the cell painting assay was performed and a morphology score was computed for each compound’s ability to revert morphology of mutant cell lines toward that of parental cells. Results of the primary morphology screen identified 58 compounds that had positive effects in two or three cell lines (Figure 6B-C). Because CDG/CDDG cell lines demonstrate elevated autophagy levels, primary screening hits were subsequently assessed for their ability to modulate autophagy by immunocytochemistry with LC3 as the marker. Twelve candidate compounds reduced autophagy in all three cell lines (Group I, Supplementary Table 2), and 10 additional candidate compounds (confirmed in at least 2 cell lines) that had a minimal, or no effect on autophagy (Group II, Supplementary Table 2).

**Figure 6.**
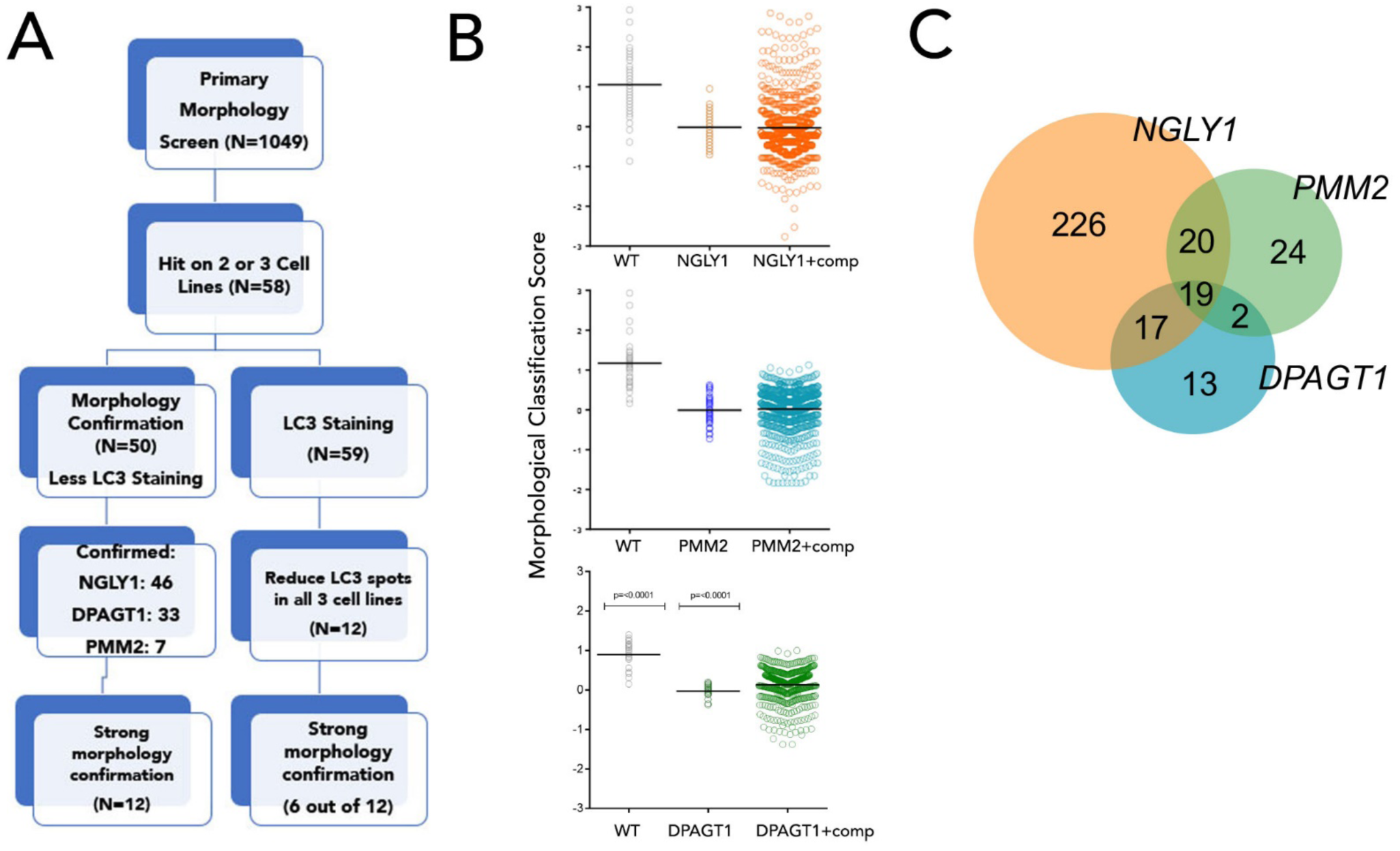
Primary screen identified compounds able to revert aberrant CDG and CDDG morphology phenotype. A, Screening workflow for selection of candidate compounds. B, Representative scatter plots of primary screen results for each genotype. Each dot represents one well. For RPE-1 and vehicle treated isogenic CDG and CDDG cell lines, N=56 replicate wells. For compound treatments, each compound was tested in N=1 well. C, Venn diagram comparing the number and overlap of compounds affecting each CDG/CDDG phenotype.

### Evaluation of compounds for amelioration of ER stress and proliferation defects

Top six candidates from each Group I and Group II were further evaluated their ability to alleviate ER stress in the CDG and CDDG lines. Additionally, compounds that improved phenotypes in all three genetic lines were added to the validation list. In order to validate the efficiency and potency of the primary screen, ten non-active compounds in the cell painting assay were selected for comparison to candidate compounds (Table 1). In total, sixteen protective compounds and 10 inactive compounds (Table 1, Supplementary Table 4) were tested for their effects on cell proliferation and s*XBP1* expression. We focused on s*XBP1* expression because it was the only ER stress marker dysregulated across all three models and the only significant ER stress marker in *PMM2*^F119L/-^ (Figure 2C-D). Dose response curves on RPE-1 cells identified the lowest non-toxic concentration for candidate compounds (see Methods).

None of the ten non-active control compounds had effects on s*XBP1* expression (Figure 7A-C, grey) or proliferation (Figure 7D-F, grey). In contrast, all active compounds impacted one or both assays in at least one of the cell lines. Compound effects appeared to be mechanism and cell line-dependent in the proliferation and *sXBP1* assays. For example, the autophagy inducing compounds (Group II) were more efficient in the CDG lines, while Group I compounds showed effects on all lines (Figure 7 and Table 1). Compounds 1, 3, 4, 5, and 14 decreased s*XBP1* expression in CDDG lines compared to DMSO treated controls (dark grey) but did not effect proliferation (Figure 7A and D). Similarly, there was no correlation between reduction of s*XBP1* expression and repair of proliferation for compounds 3, 4, 5 and 14 in CDG cell lines. Group I candidates 6 and 9 and Group II candidates 15, 25 and 26 all effectively reduced s*XBP1* expression (Figure 7A-C), and showed restoration of proliferation in CDG and CDDG lines, (Figure 7D-F). Active compounds, but not non-active controls, were able to revert aberrant cellular morphology similar to that of vehicle treated controls (Supplementary Figure 4). Together, these data validate the effectiveness of the screen, and identify sets of compounds that are able to correct aberrant cellular phenotypes associated with CDDG and CDG genotypes.

**Figure 7.**
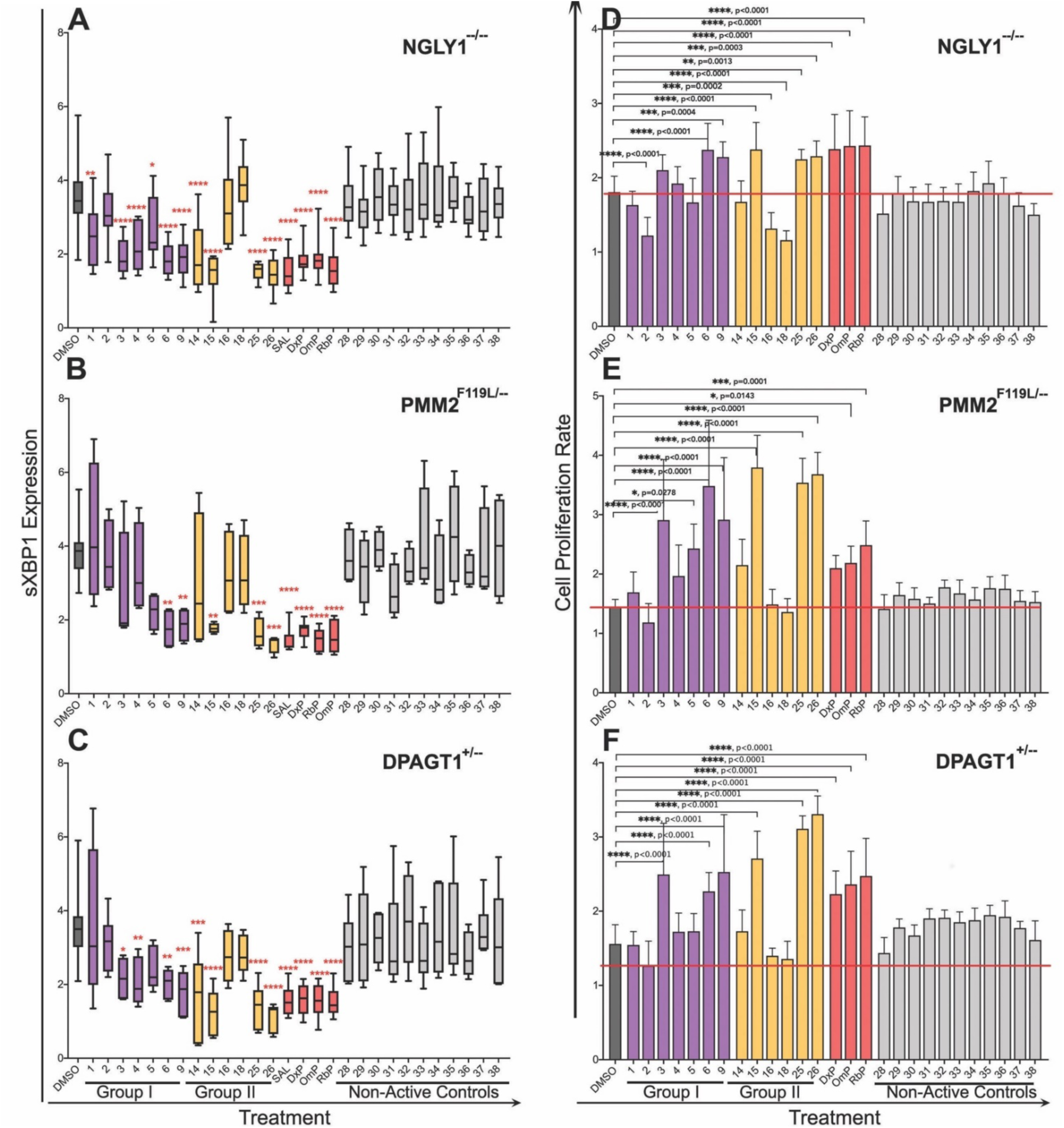
Select active compounds are able to correct proliferation and/or ER stress phenotypes in the CDG and CDDG lines. CDG and CDDG cell lines were treated with optimized concentrations of candidate and non-active control compounds and expression of s*XBP1* and cell proliferation were assessed. A-C, Expression levels of s*XBP1* in *NGLY1*^*-/-*^ (A), *PMM2*^*F119L/-*^ (B), and *DPAGT1*^*+/-*^ (C) clones treated with candidates, proton pump inhibitors (PPI; DxP, OmP, RbB) or non-active compounds. Results are presented as a ratio of s*XBP1* levels in compound-treated cells to vehicle treated cells. D-F, Modulation of proliferation rate of *NGLY1*^*-/-*^ (D), *PMM2*^*F119L/-*^ (E), and *DPAGT1*^*+/-*^ (F) lines treated with candidates, PPIs, or non-active compounds. To define cell proliferation rate, ratio of OD_590_ at 72 hrs to OD_590_ at 24 hrs post seeding (before treatment) was calculated. Data presented are an average for all clones available for a specific mutation. **, P<0.01, ***, P<0.001, ****, P<0.0001

Proton pump inhibitors (PPIs) are a class of drugs that act mainly as irreversible inhibitors of the H^+^/K^+^-ATPase pump and have been suggested as potential therapeutics for CDDG (*26*). We tested whether three PPIs (omeprazole (OmP), rabeprazole (RbP), dexlansoprazole (DxP)) could alter the cell proliferation deficits and abnormal s*XBP1* expression. We found that all of the PPIs were able to alleviate s*XBP1* expression (Figure 7A-C, red), and promote proliferation of CDG and CDDG lines (Figure 7D-F, red). Interestingly, the CDG lines were more responsive to PPIs than the CDDG line.

## Discussion

A central challenge in the development of novel therapies is the development of screenable models that focus on disease relevant phenotypes. Screens based on mutation-induced phenotypes, such as morphological differences, allows one to establish a screening assay without a full understanding of the molecular mechanisms that drive disease pathology. This creates an opportunity for the identification of new therapeutic targets as well as uncovering new insights related to etiology.

The objective of the high-content, phenotypic screen described here was to rapidly identify small molecules capable of alleviating ER stress in cellular models of monogenic disease. The rationale for our screen is that ER stress responses should be applicable across a variety of cell types, and drugs capable of alleviating ER stress will help treat symptoms of disease. Backed by the growing body of evidence linking ER stress to multiple neurological conditions and to CDG and CDDG, we reasoned that using ER stress markers as a functional readout combined with cellular phenotypes can serve as a proxy for overall cellular health on a disease background. It was important to develop the CDDG and CDG models in a cell type with uniform morphology that permits rapid and easily quantifiable morphology changes. We note that a screen in a more disease relevant cell type such as hiPSC-derived neurons may be more applicable; however, such approaches have a number of drawbacks that our approach addresses. For instance, common neuronal differentiation methods yield a heterogeneous population of cells with differing levels of maturity and morphology that renders potential molecular or morphological phenotypes difficult to identify or interpret. Moreover, the labor intensiveness and cost of differentiation methods often makes large-scale screens prohibitive. Rather, a multi-tiered strategy whereby large screens are performed on genetic cellular models with high confidence phenotypes and lead compounds are then validated in more relevant cellular and/or animal models is more efficacious.

A majority of the active compounds in our screen, #s 3, 4, 6, 9, 15, 25, are reported to affect microtubules (*27-40*) (Supplementary Table 3) either through direct effects on microtubules themselves or by targeting proteins (e.g. kinases) that regulate microtubule dynamics. Multiple compounds converging on the regulation of microtubules lends support to their involvement in ER stress responses (*41*). Furthermore, compounds, structurally similar to compound 6, 9, 15, 25 and 26 (see Supplementary Table 4 for structures) are reported to prevent ER stress in multiple cellular systems through inhibition of NADPH oxidase 2 (NOX2) (*42*), isocitrate dehydrogenase 1 (IDH1) (*43, 44*), various kinases, such as CDK (*45-48*), JAK1/2 kinases and STAT signaling (*49-51*), general control nonderepressible 2 (GSN2) kinase (*52, 53*), and growth factor receptor kinases (*54-57*) (Supplementary Table 3). CDG Type Ia (PMM2-CDG) patients often present with cerebellar atrophy (*58*), that has been attributed to defective ER stress response (*8*).

Our study describes, to our knowledge, the first example of a high-throughput screen on genetically modified human cells for three monogenic diseases with a shared endogenous molecular phenotype. Here we focused on a biological process, ER stress, thought to unite a number of genetic and more common diseases, and successfully identified bioactive ER stress diminishing compounds through unbiased morphological screening. This work has shown it is possible to develop cellular models for CDGs that possess screenable phenotypes able to identify compounds that alleviate molecular and morphological phenotypes caused by these genetic conditions, thereby establishing a platform to identify targeted and common treatments for CDDG and CDGs. Due to the genetic heterogeneity of CDGs, it will be important to apply similar analyses to other genetic causes of CDG to determine whether there are compounds that alleviate the ER stress-related symptoms across a wide range of disease. Beyond establishing a paradigm for identifying therapeutic compounds for rare monogenic diseases, this work suggests a direction for identifying compounds able to alleviate the symptoms related to ER stress in more common diseases characterized by ER stress including neurodegenerative diseases.

Loss of microtubule mass or altered microtubule dynamics in axons and dendrites are major contributors to neurodegenerative diseases such as ALS, Parkinson’s disease, Alzheimer’s disease, and several tauopathies (*59*). A recent screen of a ∼100 small molecules identified compounds protective of ER stress in a chemically-induced ER stress model of ALS (SOD1^G93A^) neurodegeneration (*60*). Future studies will determine whether compounds that affect microtubule dynamics are able to prevent disease-relevant phenotypes in cellular models of neurodegenerative diseases.

## Materials and Methods

### CRISPR/Cas9 Genome Editing of hTERT RPE-1 Cells

CDG and CDDG lines were generated by CRISPR/Cas9 genome editing of hTERT RPE-1 (ATCC, CRL-4000TM) at the Columbia Stem Cell Core Facility. Promoter (U6) and gRNA scaffolds were synthesized by Integrated DNA Technologies (IDT) and cloned into the pCR-Blunt II-TOPO plasmid (ThermoFisher Scientific, cat. K280002). Nucleofector (Lonza) was employed to introduce gRNA and Cas9-GFP plasmids into hTERT RPE-1 cells. After nucleofection, single colonies were manually picked into either 96-well plates or 10 cm dishes. Cells were incubated for ten days to reach confluency in a 96-well plate or visible colonies in a 10 cm cell culture dish. For each colony, DNA was extracted by the KAPA Mouse Genotyping Kit (KAPA Biosystems) and genotyped by Sanger sequencing.

Guide RNA scaffold and termination signal: GTTTTAGAGCTAGAAATAGCAAGTTAAAATAAGGCTAGTCCGTTATCAACTTGAAAAAGTGGCACCGAGTCGGTGCTTTTTTT

gRNA *NGLY1:* GGTGATTGCCAGAAGAACTAAGG, *PMM2:* GAATTCAATGAAAGTACCCCTGG, *DPAGT1:* CATGATCTTCCTGGGCTTTGCGG

### Chemicals

Tunicamycin (cat. 3516), salubrinal (cat. 2347), omeprazole (cat. 2583) were purchased from Tocris. Dexlansoprazole (cat. HY-13662B) and rapamycin (cat. HY-10219) were obtained from MedChemExpress. Rabeprazole (cat. 14939) was from Cayman Chemicals. All screened compounds were provided by Janssen Pharmaceuticals. All compounds were suspended in DMSO, aliquoted, and stored at −20°C.

### Proliferation measurements, MTT assay

Cells were seeded in 96-well tissue culture plates (1× 10^3^ cells per well) and treated the next day as described in ‘Results’. At the indicated time points, the medium was removed, and fresh medium containing 0.5 mg/ml MTT (3-(4,5-dimethylthiazol-2-yl)-2,5-diphenyltetrazolium bromide, Sigma-Aldrich) was added to each well. The cells were incubated at 37°C for 4 h and then an equal volume of solubilization solution (0.01 N HCl in 10% SDS) was added to each well and mixed thoroughly. The optical density from the plates was read at 570 nm, and background absorbance read at 690 nm. A statistical analysis of the results was performed using the GraphPad Prism software (v.8.2.0). One-way ANOVA multiple comparisons and Dunnett test were used to determine the equality of the means of different samples. The confidence level (p) was 0.05.

### Quantitative RT-PCR

Total RNA was extracted by RNeasy Plus Mini kit (QIAGEN, cat. 74136). Total RNA (500 ng) was reverse-transcribed with random primers using Superscript IV Reverse Transcriptase kit (ThermoFisher Scientific, cat. 18091200). One µL of cDNA was used in each qPCR reaction on a QuantStudio 5 (ThermoFisher Scientific) using SYBR Green PCR Master Mix (ThermoFisher Scientific, cat. 4364344). PCR primers detecting spliced and unspliced *XBP1* expression were as described (*61, 62*). Each reaction was performed in duplicate with an initial holding step of 95°C (10 min) followed by 40 cycles of 95°C (10 s) and 55°C (30 s). N = ≥3 for each experiment. The relative expression levels of target genes were normalized to that of the reference *GAPDH* gene by using the ΔΔCt method (*63*). The fold change in expression for each sample is relative to parental hTERT RPE-1 cells treated with vehicle. Data were graphed using Prism 8 software. The following sets of primers were used for real-time PCR: for human total *XBP1*, TGGCCGGGTCTGCTGAGTCCG and ATCCATGGGAAGATGTTCTGG; spliced *XBP1*, CTGAGTCCGAATCAGGTGCAG and ATCCATGGGAAGATGTTCTGG; for human *GAPDH*, ACAGTCAGCCGCATCTTCTT and TTGATTTTGGAGGGATCTCG.

### Immunoblot analysis

Cell lysates were prepared on ice in RIPA buffer (Sigma-Aldrich, cat. R0278) with freshly added cocktail of proteases (Millipore-Sigma, cat. 11836170001) and phosphatases (Millipore Sigma, cat.4906837001) inhibitors. Protein concentrations were determined using Pierce BCA Protein assay kit (ThermoFisher Scientific, cat.23227). Whole cell lysates (20–50 μg) were subjected to SDS-PAGE, transferred to PVDF membrane (Immobilon-P, Millipore, cat. IPVH00010), and membranes were blocked and probed by overnight incubation with appropriate primary antibodies. Antibodies: p-eIF2α (Ser51) and eIF-2α (Cell Signaling Technology, cat. 9721 and 9722). NGLY1 rabbit polyclonal antibody (Bethyl Laboratories, cat. A305-547A-T), mouse monoclonal ATF6 and mouse monoclonal PMM2 (2E9) (Novus Biologicals, cat. NBP1-40256 and H00005373-M01). DPAGT1 rabbit polyclonal antibody (Abcam, cat. ab116667). Bound antibodies were visualized with corresponding HRP-conjugated secondary antibodies (ThermoFisher Scientific, cat. 32260 and 31430) and detected by ECL chemistry (SuperPico West, ThermoFisher Scientific, cat. 34580). Western blots were quantitatively analyzed via laser-scanning densitometry using NIH ImageJ Version 1.52k software. IB for β-actin (Santa Cruz, cat. sc-47778) was used to ascertain equal protein loading across samples.

### Immunocytochemistry

hTERT RPE-1 and the isogenic mutant lines were seeded on poly-D-lysine-coated 12 mm glass coverslips at a density of 5×10^4^ cells/well. Next day, cells were quickly washed with phosphate buffered saline (PBS) and then fixed with 4% paraformaldehyde (PFA) for 10 min at RT. Following three washes in PBS, cells were incubated with permeabilization buffer (1% Triton X-100 in PBS) for 15 min at RT, and block in staining buffer (1% bovine serum albumin, 0.1% Triton X-100 in PBS) for 1 hour at RT. Cells were incubated with primary antibodies diluted in staining buffer for 1.5 hours at RT, washed three times with PBS, then incubated with the appropriate fluorescently labeled secondary antibodies diluted in staining solution for 30 min at RT in the dark. Finally, the cells were washed three times with PBS, and coverslips were mounted on microscopy slides using Prolong Antifade DAPI (Invitrogen) and allowed to cure overnight at RT in the dark before imaging. Primary antibodies: mouse monoclonal PMM2 (2E9) (Novus Biologicals, H00005373-M01, 1:500), DPAGT1 rabbit polyclonal antibody (Abcam, cat. ab116667, 1:500), mouse anti-CHOP clone L637F (Cell Signaling Technology, cat. 2895, 1:3200), rabbit anti-GADD153/CHOP (Novus Biologicals, cat. NBP2-58505, 1:500), mouse anti-LAMP-1/CD107a, clone H4A3 (Novus Biologicals, cat. NBP225183, 1:500), mouse anti-SQSTM1/p62 (Abcam, cat. ab56416, 1:200), rabbit anti-P4HB/PDIA1 (Abcam, cat. ab3672, 1:200), LC3A (D50G8) XP rabbit monoclonal antibody (Cell Signaling Technology, cat. 4599). Secondary antibodies, donkey anti-mouse or donkey anti-rabbit conjugated to either Alexa Fluor 488 or 568 (Invitrogen, cat. A32766, A32790, A10042, A10037, 1:1000). Imaging was performed on an inverted Zeiss AxioObserver Z1 fluorescent microscope equipped with an AxioCam 503 mono camera and filters for 405 nm, 488 nm, and 568 nm. Images were acquired with Zen 2 software and post-processing was performed with Zen 2 and AdobePhotoshop.

### Senescence detection, β-Galactosidase Staining

Cells were seeded in 6 well plates (1×10^5^/well) overnight, and next day were stained for β-galactosidase using Senescence β-Galactosidase Staining Kit (Cell Signaling Technology, cat. 9860) according to manufacturer’s instruction. The images were acquired, and the number of stained cells was counted using Zeiss Primovert inverted brightfield and phase contrast microscope equipped with AxioCam ERc5s camera.

### Apoptosis detection by Annexin-V binding assay

Cells were suspended in 1x Annexin V binding buffer (10 mM HEPES/NaOH, pH 7.4, 150 mM NaCl, 2.5 mM CaCl_2_) before staining with APC-labeled Annexin-V (BD Biosciences, cat. 550474) according to the manufacturer’s instructions. Propidium iodide (PI) was added to the samples after staining with Annexin-V to exclude late apoptotic and necrotic cells. FACS was performed immediately after staining on FACSCelesta (BD Biosystems). Data were analyzed using FlowJo v. 10.5.3 and Prism8 v8.2.0 software.

### Autophagy detection

using CYTO-ID® Autophagy detection kit (ENZO Life Sciences, cat. ENZ-51031-K200). For fluorescent microscopy, cells were seeded on poly-D-lysine-coated 12 mm glass coverslips at a density of 5×10^4^ cells/well. The next day, cells were washed with PBS, stained with CYTO-ID ® Green reagent according to manufacturer’s instructions, and observed using an inverted Zeiss AxioObserver Z1 epifluorescent microscope equipped with an AxioCam 503 mono camera. Image post-processing was performed with Zen 2 and Adobe Photoshop software. For flow cytometry analysis, cells were trypsinized, collected by centrifugation, washed with PBS and stained using CYTO-ID ® stain solution (ENZO Life Sciences, cat. ENZ-51031-K200) according to manufacturer’s instructions and immediately analyzed on a FACSCelesta cytometer (BD Biosystems). Data were analyzed using FlowJo v. 10.5.3 and Prism8 v8.2.0 software.

### Autophagy detection by p62 and LAMP1 staining

Cells were collected by trypsinization, washed with PBS and fixed with 4% PFA/PBS for 15 min at RT. Fixed cells (10^6^ cells/sample) were permeabilized using Intracellular Staining Permeabilization Wash buffer (BioLegend, cat.421002) according to the manufacturer instructions and stained with primary mouse monoclonal antibodies for SQSTM1/p62 (Abcam, cat. ab56416, 1:1000), LAMP1/CD107a, clone H4A3 (Novus Biologicals, cat. NBP225183, 1:500), or mouse IgG isotype control antibody (ThermoFisher Scientific, cat. MA5-14453, 1:500) for 1 hr at RT. After two washes with PBS, cells were incubated for 30 min at RT with Alexa-488 labeled secondary goat anti-mouse antibody (ThermoFisher Scientific, cat. A32723, 1:1000), washed again twice with PBS and analyzed using flow cytometry on a FACSCelesta cytometer (BD Biosystems). The data were processed using FlowJo v. 10.5.3 and Prism8 v8.2.0 software.

### Cellular morphology assessment by immunostaining

Cells were seeded in 96 well plates at a density of 1×10^3^ cells/well, and next day were treated with tested compounds, vehicle (DMSO) or positive controls as described in Materials and Methods. After 24 hrs, cells were washed with PBS, fixed with 4% PFA/PBS for 15 min at RT, blocked with 1% BSA/PBS for 30 min and stained with Alexa 568 labeled phalloidin (ThermoFisher Scientific, cat. A12380) for 30 min. After two washes with PBS, cells were stained with 300nM DAPI (BD Pharmingen, cat. 564907). Imaging was performed with an inverted Zeiss AxioObserver Z1 epifluorescent microscope equipped with an AxioCam 503 mono camera, and images acquired with the Zen 2 software. Post-processing was performed with Zen 2 and AdobePhotoshop software.

### High-content imaging and compound screening

The hTERT RPE-1 cells and *NGLY1*^*-/-*^, *PMM2*^*F119L/-*^, *and DPAGT1*^*+/-*^ mutant lines were plated in 384 well plates at a density of 3,000 cells per well. The next day compounds were added to the cells at a final concentration of 10 µM and incubated for 24 hours. MitoTracker Red (Molecular Probes, cat. M7512) mitochondrial stain was added to the media, following the manufacturer’s protocol. After 30 minutes of labeling, media was removed, cells were fixed in 4% PFA in PBS, washed, permeabilized with 0.1% NP-40, and blocked with 3% BSA in PBS overnight. For staining, ConcanavalinA-488 (Molecular Probes, cat. C11252), phalloidin-547 (Molecular Probes, cat. A22283), and DAPI were added to the wells, then washed before imaging. Images were acquired on a Molecular Devices Image Express microscope at 4 fields per well. Feature extraction from images was done with Perkin-Elmer Columbus Image Analysis software, and feature analysis and hit determination was performed using TIBCO Spotfire analysis package.

### Secondary validation of selected compounds

Candidate and control compounds from the high-throughput screen were validated in using two assays, MTT to assess proliferation and RT-qPCR for s*XBP1* expression as described above. Parental hTERT RPE-1 cells and their edited clones were seeded in 6-well (10^5^ cells/well, for RNA assay) or 96-well (10^3^ cells/well, for MTT assay) plates and in a humidified incubator at 37°C and 5% CO_2_, and treated with candidate or control compounds. Dose response curves on hTERT RPE-1 cells identified the lowest non-toxic concentration for each of the candidate compounds. Cells were treated for 24 h with Group I (5.0 nM), Group II (1.0 nM), or non-active control (10 µM) compounds. Concentrations of PPI were; DxP 50 µM, OmP and RbP 100 µM. Post treatment, total RNA was collected in RLT buffer (QIAGEN) containing β-mercaptoethanol for RT-qPCR. Alternatively, cells were subjected to MTT assay at days 0, 1, 3 and 5 post-treatment.

### Statistical analyses

Results are expressed as mean ± SEM for a minimum of 3 independent experiments. Sample size and statistical tests are detailed in the figure legends. Statistical analysis was performed using one-way ANOVA followed by Dunnett multiple comparisons post-test to compare each condition to vehicle-treated controls. P values ≤ .05 were considered significant.

## Acknowledgments

We thank Guy Ludwig and other past and present members of the Institute for Genomic Medicine for insights and helpful discussions.

## Author contributions

(according to CRediT, https://www.casrai.org/credit.html)

**Conceptualization:** D.B.G., M.J.B., Y-F.L., M.V.W., J.W.

**Data Curation:** I.V.L., M.V.W., S.S.

**Formal Analysis:** I.V.L., M.V.W., S.S.

**Investigation:** I.V.L., M.V.W., S.S.

**Methodology:** I.V.L., M.V.W., Y-F.L.

**Project Administration:** I.V.L., M.V.W., A.A-O., J.W., M.J.B.

**Resources:**

**Software:**

**Supervision:** M.J.B., J.W., D.B.G.

**Validation:** I.V.L., M.V.W., S.S., M.J.B.

**Visualization:** I.V.L., M.J.B.

**Writing – Original Draft:** I.V.L., M.J.B., D.B.G.

**Writing – Review & Editing:** I.V.L., M.V.W., M.B.H., M.J.B., D.B.G.

