## Supplementary Data for "Precision genetic cellular models identify therapies protective against endoplasmic reticulum stress"

**List of Supplementary Materials**

4 Supplementary Figures:

**Supplementary Figure 1.** Genotype of CDG and CDDG RPE-1 lines.

**Supplementary Figure 2.** Assessment of senescence and apoptosis.

**Supplementary Figure 3.** Assessment of autophagy by flow cytometry.

**Supplementary Figure 4.** Reversion of cellular morphology by candidate compounds

4 Supplementary Tables**:**

**Supplementary Table 1.** Summary of genome edited CDG and CDDG cell line genotypes.

**Supplementary Table 2.** A list of “hit” compounds determined by HTS cell painting assay and nominated for further biochemical testing

**Supplementary Table 3.** Chemical names and properties of compounds with confirmed biological activity.

**Supplementary Table 4.** Predicted structures for compounds with confirmed biological activity.

**Supplementary Figure 1**

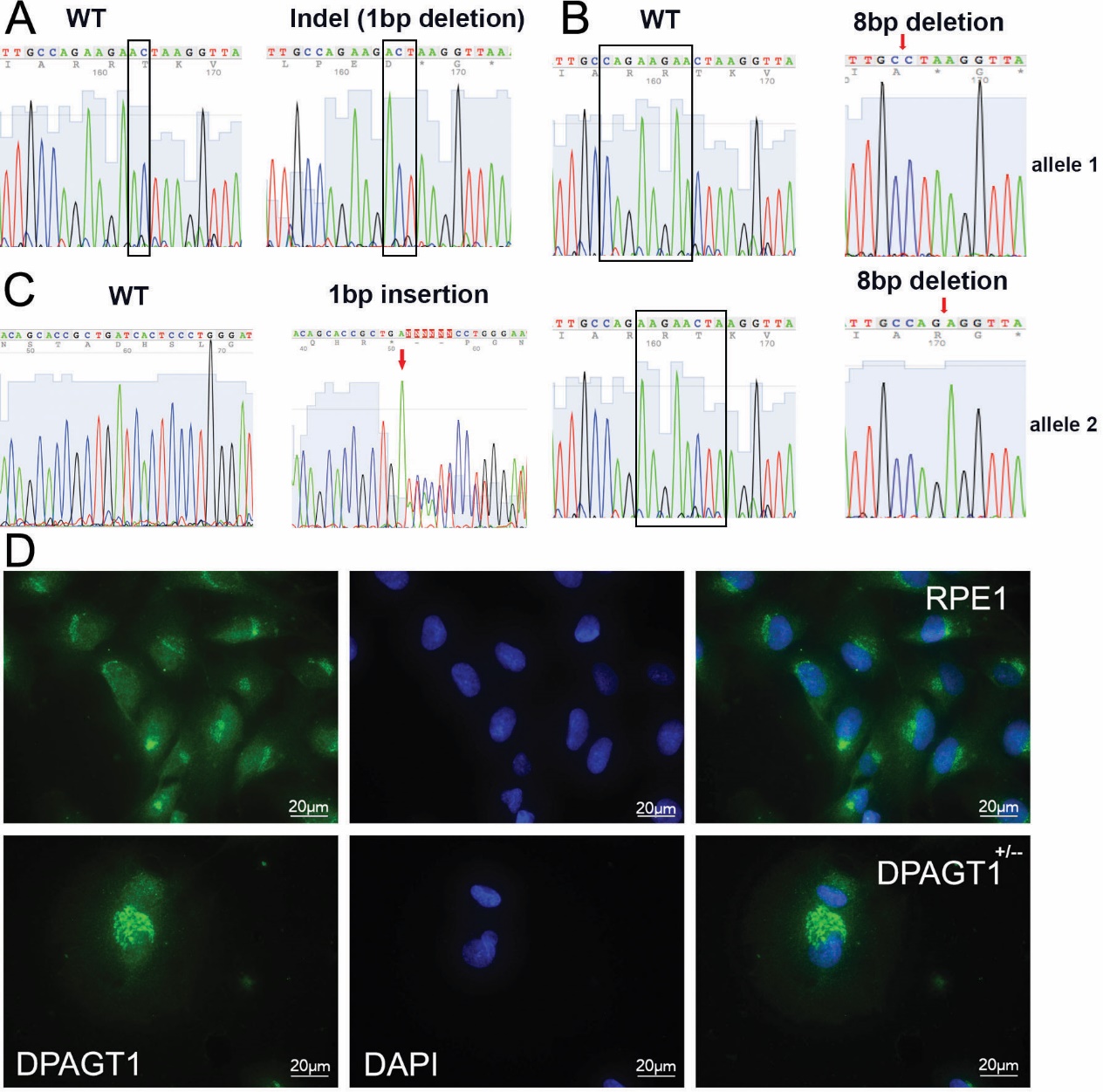

**Supplementary Figure 1.** **Genotype of CDG and CDDG RPE-1 lines.**

A-C, Electropherogram traces for parental RPE-1 cells and CDDG, *NGLY1*^-/-^ D3 (A) and *NGLY1*^-/-^ E6 (B), and CDG *DPAGT1*^+/-^ D12 (C) lines. B and C, Immunoblot images for target proteins in RPE-1 and isogenic CDG and CDDG lines. D, Representative immunofluorescence images for staining for DPAGT1 protein in parental RPE-1 and CDG *DPAGT1*^+/-^ D12 lines.

**Supplementary Figure 2**

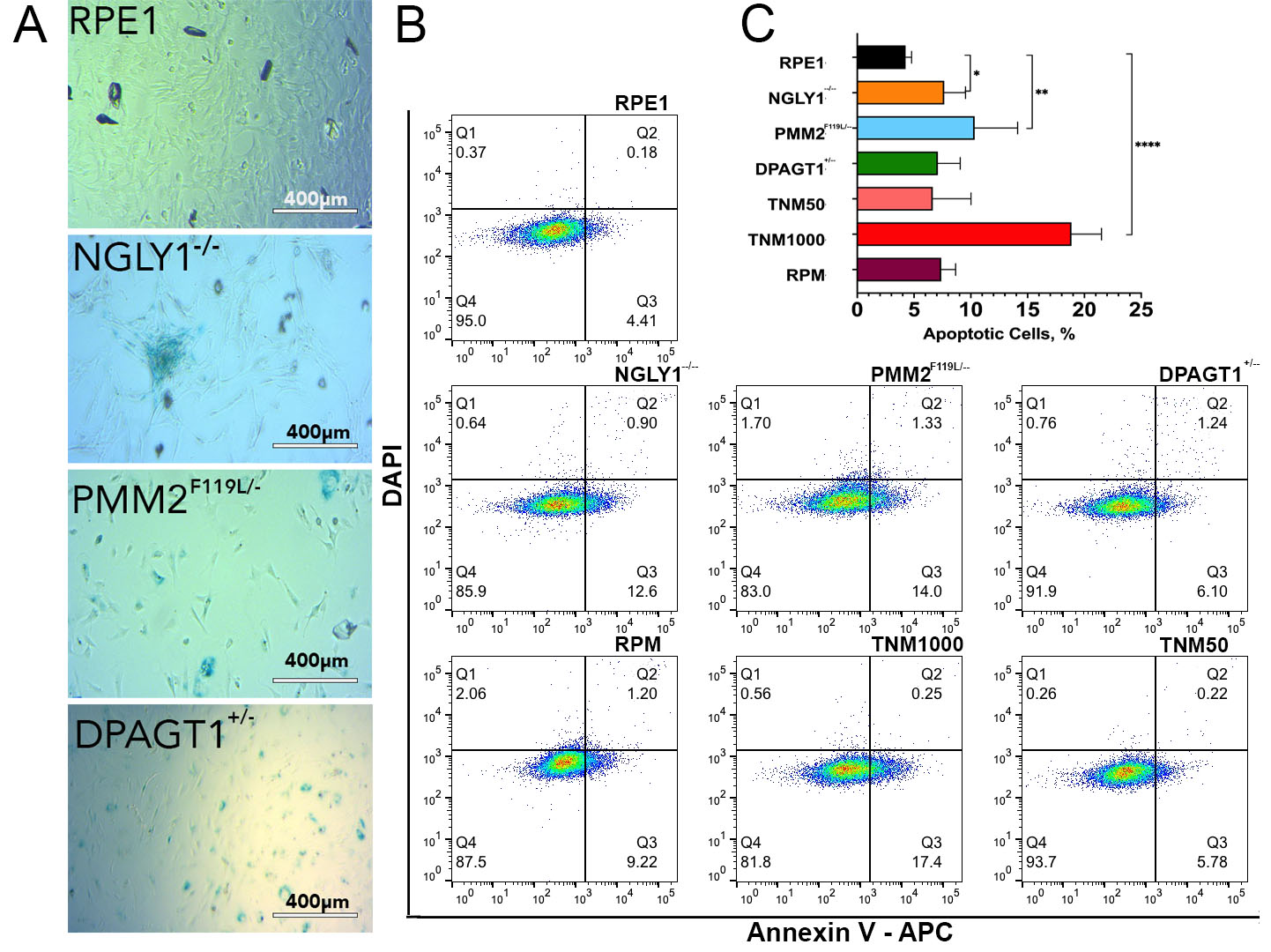

**Supplementary Figure 2. Assessment of senescence and apoptosis.** A, Representative images of β-galactosidase senescence staining of parental RPE-1 and CDDG, *NGLY1*^-/-^ D11, and CDG *PMM2* ^F119L/-^ A3 and *DPAGT1*^+/-^ D5 lines. B. Representative dot-blots of apoptosis detection (Annexin V staining) in RPE-1, CDG and CDDG lines. Tunicamycin was used as a positive control for apoptosis (1.0 µM, 24 h) and chronic ER stress (0.05 µM, 24 h). Rapamycin treatment (RPM, 500nM) was used as a control for autophagy induction. C, Quantification of apoptosis by flow cytometry (N=3 experiments). *, P<0.05, **, P<0.01, ****, P<0.0001.

**Supplementary Figure 3**

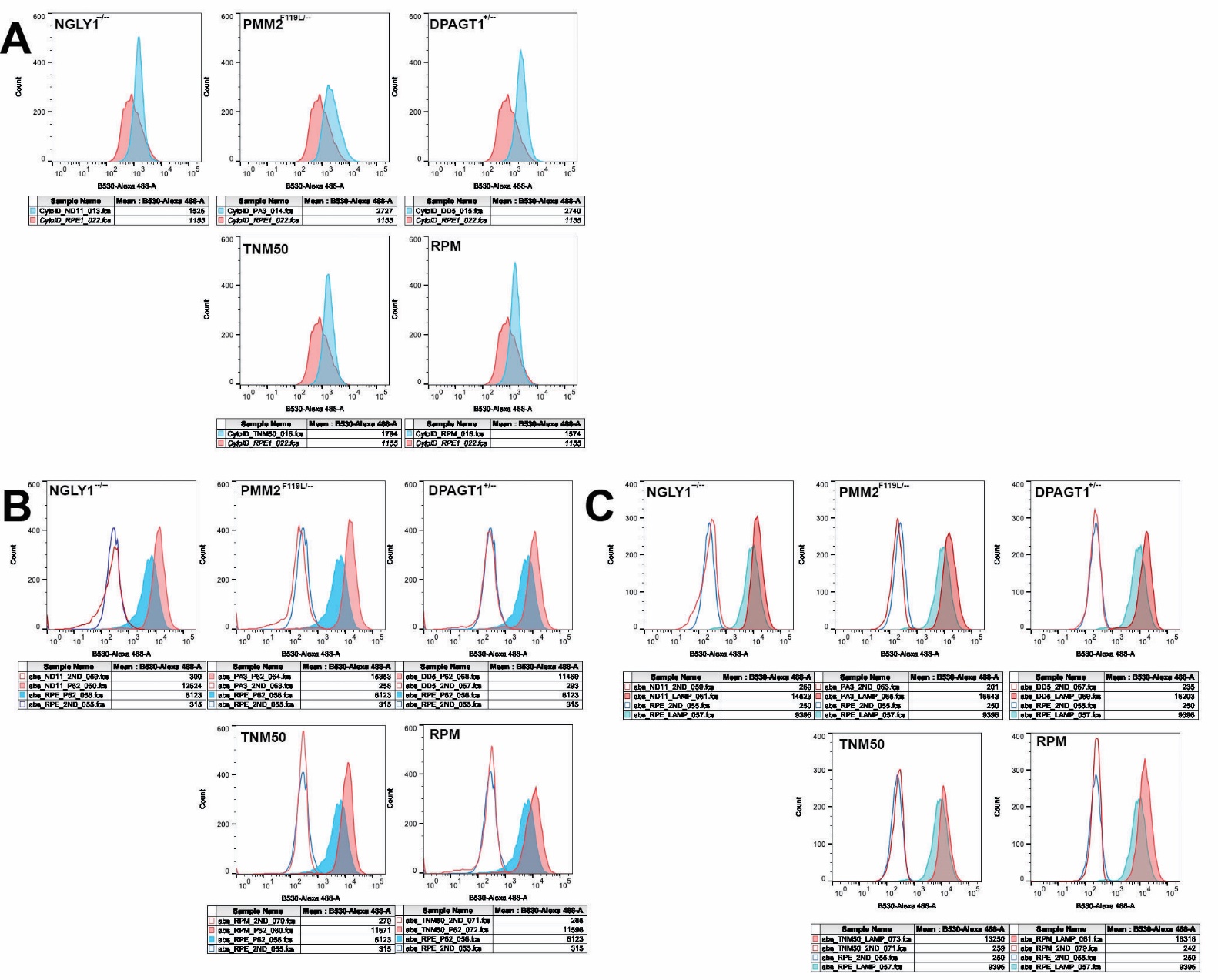

**Supplementary Figure 3. Assessment of autophagy by flow cytometry.** Representative histogram overlays of RPE-1 and isogenic CDG and CDDG cell lines stained with Cyto-ID® reagent (A), anti-p62/SQATM1 (B) and anti-LAMP1 (C) antibodies. Median Fluorescence Intensity (MFI) values for each sample are indicated in the tables below the images.

**Supplementary Figure 4**

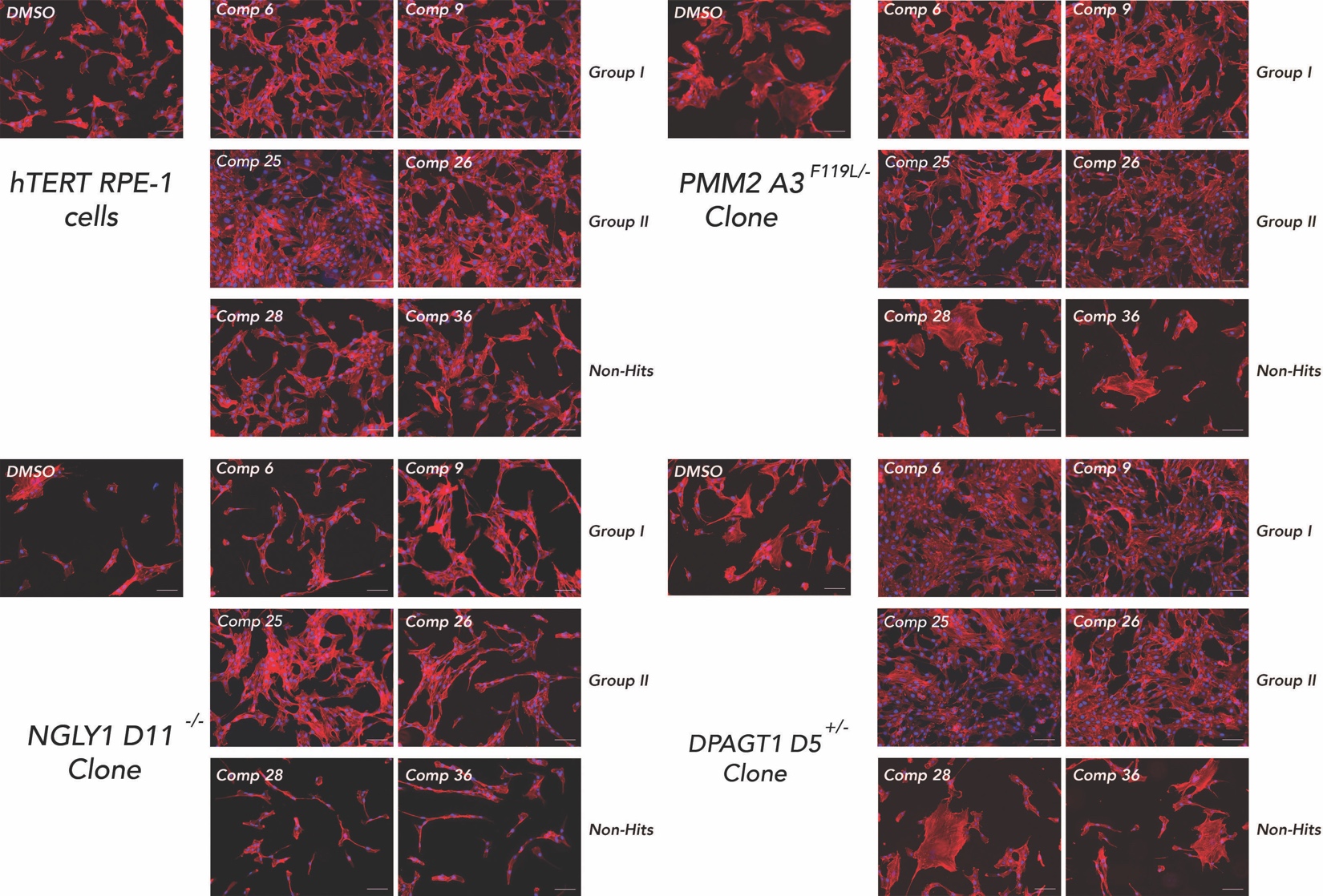

**Supplementary Figure 4.** **Reversion of cellular morphology by candidate compounds.** Representative images of morphological changes in parental RPE-1 and isogenic CDDG, *NGLY1*^-/-^ D11, and CDG *PMM2* ^F119L/-^ A3 and *DPAGT1*^+/-^ D5 lines upon treatment with indicated compounds.

**Supplementary Table 1**

Summary of genome edited CDG and CDDG cell line genotypes. All genotypes were confirmed by Sanger sequencing.

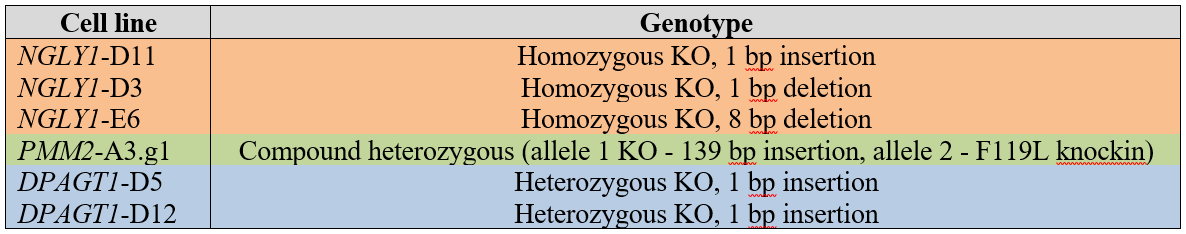

**Supplementary Table 2.**

A list of “hit” compounds determined by HTS cell painting assay and nominated for further biochemical testing

|  | Primary Screen Hit (n=1) | | | Confirmation (n=4) | | | LC3 puncta (% control) | | |
| --- | --- | --- | --- | --- | --- | --- | --- | --- | --- |
| Index | *DPAGT1* | *NGLY1* | *PMM2* | *DPAGT1* | *NGLY1* | *PMM2* | *DPAGT1* | *NGLY1* | *PMM2* |
| 1 | 1 | 1 | 1 | 4 | 4 | 1 | 31 | 30 | 23 |
| 2 |  | 1 | 1 | 4 | 2 |  | 22 | 11 | 8 |
| 3 | 1 | 1 | 1 | 3 | 4 |  | 27 | 13 | 11 |
| 4 | 1 | 1 | 1 | 3 | 3 |  | 27 | 18 | 18 |
| 5 |  | 1 | 1 | 2 | 2 | 1 | 27 | 14 | 15 |
| 6 |  | 1 | 1 | 2 | 2 |  | 29 | 27 | 30 |
| 7 | 1 | 1 |  | 2 | 1 |  | 42 | 30 | 19 |
| 8 | 1 | 1 |  | 1 | 4 |  | 47 | 10 | 13 |
| 9 |  | 1 | 1 | 1 | 2 | 1 | 27 | 16 | 11 |
| 10 | 1 | 1 | 1 |  |  |  | 26 | 13 | 12 |
| 11 | 1 | 1 | 1 |  |  |  | 37 | 18 | 10 |
| 12 |  | 1 | 1 |  |  |  | 39 | 32 | 30 |
| 14 | 1 | 1 |  | 4 | 3 |  | 95 | 47 | 35 |
| 15 | 1 | 1 |  | 4 | 2 |  | 85 | 55 | 29 |
| 16 | 1 | 1 | 1 | 3 | 4 |  | 80 | 34 | 40 |
| 18 | 1 | 1 |  | 3 | 3 |  | 131 | 63 | 51 |
| 19 |  | 1 | 1 | 3 | 2 |  | 105 | 66 | 64 |
| 21 | 1 | 1 |  | 2 | 3 |  | 85 | 40 | 33 |
| 24 | 1 | 1 | 1 | 1 | 3 |  | 109 | 37 | 34 |
| 25 |  | 1 | 1 | 1 | 3 | 1 | 117 | 48 | 52 |
| 26 | 1 | 1 |  | 1 | 2 | 1 | 96 | 45 | 37 |
| 27 |  | 1 | 1 |  | 2 |  | 108 | 42 | 26 |

**Supplementary Table 3.**

Chemical Names and Properties of Compounds with Confirmed Biological Activity.

| Index | CAS Number | IUPAC Name | Name* and/or Chemical Class | Target |
| --- | --- | --- | --- | --- |
| 3 | 31430-18-9 | methyl N-[5-(thiophene-2-carbonyl)-1H-benzimidazol-2-yl]carbamate | Nocodazole  (Benzimidazoles) | Microtubules (*1-3*) |
| 4 | 14255-87-9 | methyl N-(5-butyl-1H-benzimidazol-2-yl) carbamate | Parbendazole (Benzimidazoles) | Microtubules (*4-6*) |
| 6 | 840534-89-6 | 2-[3-[[3-(3-fluorophenyl)triazolo[4,5-d]pyrimidin-5-yl] amino]phenyl]acetic acid | Triazolopyrimidines | Microtubules (*7*), NOX2 (*8*), TDP2 (*9*), GSN2, PERK, HRI, IRE1 inhibitor (*10, 11*) |
| 9 | 518-28-5 | (5R,5aR,8aR,9R)-5-hydroxy-9-(3,4,5-trimethoxyphenyl)-5a,6,8a,9-tetrahydro-5H-isobenzofuro[6,5-f][1,3]benzodioxol-8-one | Podophyllotoxin (Lignans,  Furonaphthodioxols) | Microtubules (*12, 13*), IGF 1R (*14, 15*), IDH1 (*16*), TDP1/2 (*17*) |
| 15 | 935666-88-9 | 5-chloro-N2-[(1S)-1-(5-fluoropyrimidin-2-yl)ethyl]-N4-(5-methyl-1H-pyrazol-3-yl)pyrimidine-2,4-diamine | AZD1480  (Aminopyrazoles) | Microtubules (*18*), JAK1/2 (*18-20*), ALK, LTK, FGFR, RET and TRK kinases inhibitor (*21*), STAT3 and STAT5A inhibitor (*22, 23*), IDH1 inhibitor (*19*) |
| 25 | 627517-32-2 | [3-(3-fluoroanilino)-6,7-dimethoxy-4H-indeno[1,2-c]pyrazol-1-yl]methyl butanoate | Aminopyrazoles  Benzenesulfonamides | Microtubules (*24-27*)  PDGF-R kinase inhibitor (*28*) |
| 26 | 842128-50-1 | 3-[5-[3-(2-aminopyrimidin-4-yl)anilino]triazolo[4,5-d]pyrimidin-3-yl]benzenesulfonamide | Aminopyrimidines, benzensulfonamides | Possible CDK1/2/5/9 inhibitor (*29-32*) |

*Commercial name if available

**Supplementary Table 4.**

Predicted structures for compounds with confirmed biological activity. Group I compounds (left), Group II compounds (right).

| **Index** | **CAS No.** | **GCRS Sketch** | **SMILES** |
| --- | --- | --- | --- |
| 14 | 443798-87-6 | 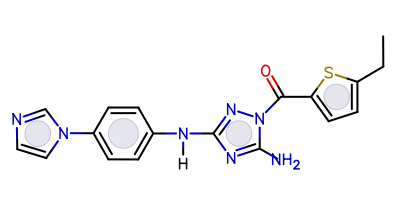 | 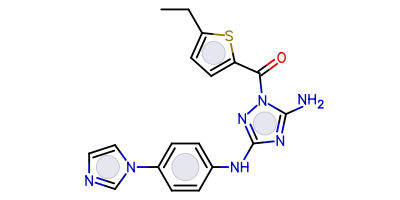 |
| 15 | 935666-88-9 | 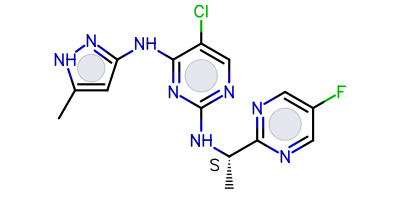 | 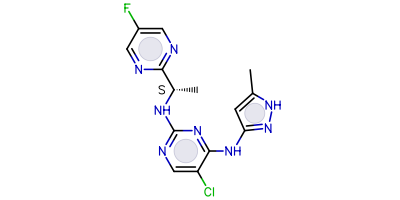 |
| 16 | 244768-00-1 | 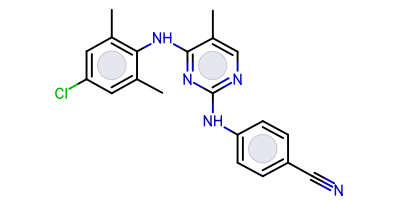 | 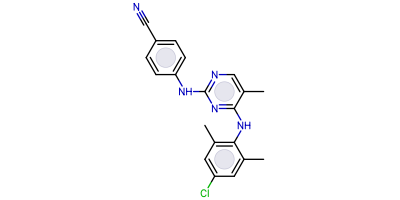 |
| 18 | 443799-16-4 | 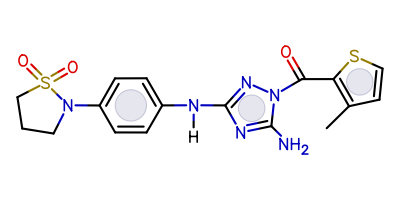 | 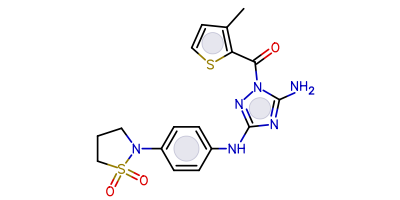 |
| 25 | 244767-84-8 | 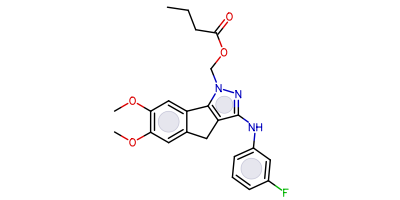 | 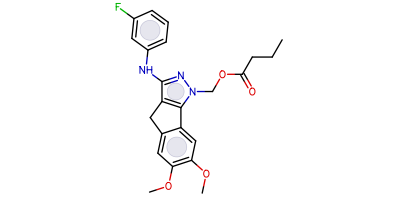 |
| 26 | 627517-32-2 | 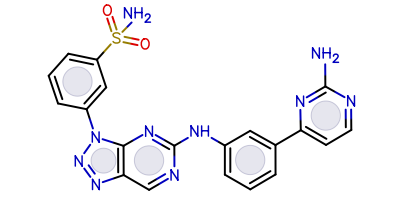 | 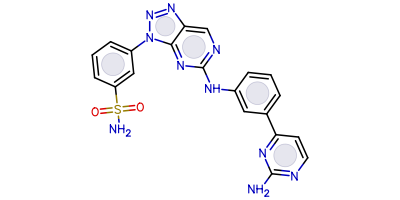 |

| **Index** | **CAS No.** | **GCRS Sketch** | **SMILES** |
| --- | --- | --- | --- |
| 1 | 1092504-43-2 | 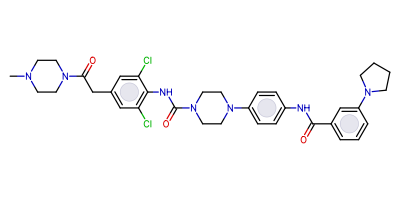 | 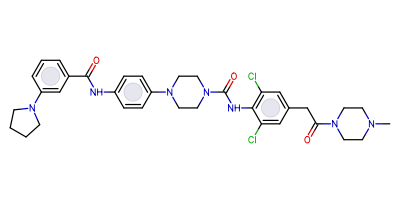 |
| 2 | 126452-70-8 | 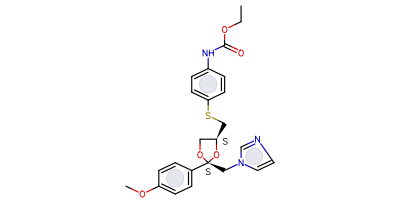 | 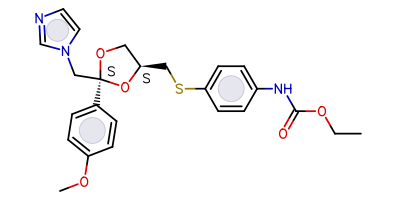 |
| 3 | 31430-18-9 | 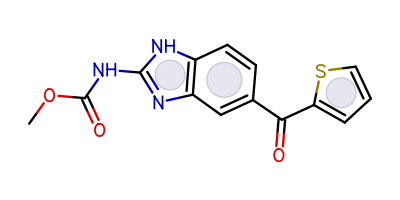 | 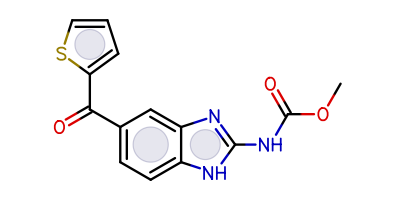 |
| 4 | 14255-87-9 | 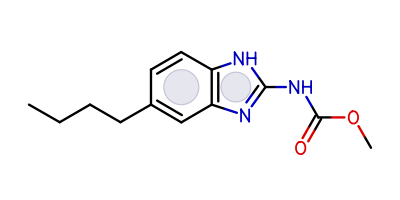 | 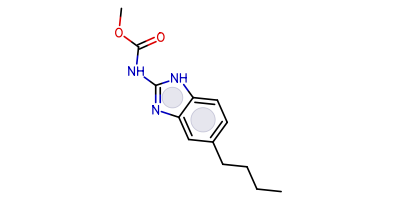 |
| 5 | 627512-62-3 | 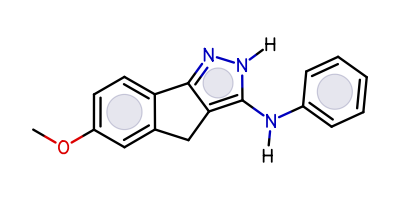 | 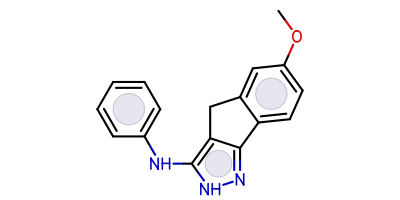 |
| 6 | 840534-89-6 | 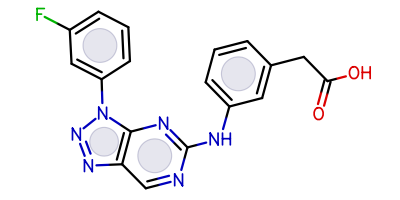 | 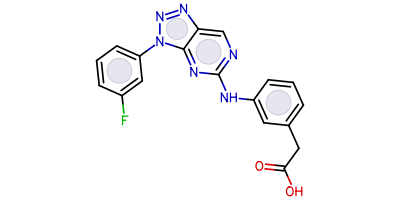 |
| 9 | 518-28-5 | 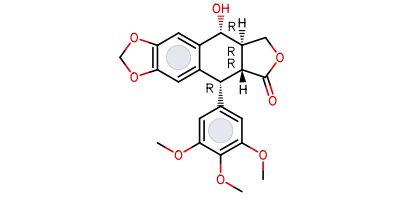 | 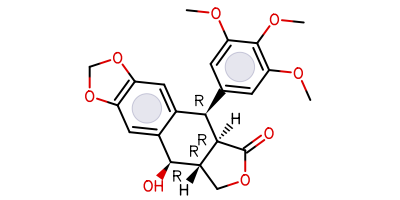 |

**Supplementary references.**

6.	A. Ganguly, H. Zhang, R. Sharma, S. Parsons, K. D. Patel, Isolation of human umbilical vein endothelial cells and their use in the study of neutrophil transmigration under flow conditions. *J Vis Exp*, e4032 (2012).
